# Absence of Sensory Cooling Activity and Cooling Agents from California’s New “Non-Menthol” Cigarettes Marketed in 2025

**DOI:** 10.1101/2025.11.17.688932

**Authors:** Sairam V. Jabba, Hanno C. Erythropel, Paul T. Anastas, Stephanie S. O’Malley, Suchitra Krishnan-Sarin, Julie B. Zimmerman, Sven E. Jordt

## Abstract

**Background:** Since late 2022, the sale of most flavoured tobacco products has been prohibited in California, including menthol cigarettes. Tobacco companies responded by introducing “non-menthol” cigarettes in which menthol was replaced with WS-3, an odorless synthetic cooling agent to elicit cooling sensations similar to menthol. Legislation enacted in 2024 banned the addition of cooling characterizing flavours in tobacco products in California. However, the industry continues to market “non-menthol” cigarettes in the state, with very similar package designs. It is unknown whether these cigarettes contain a cooling agent.

**Methods:** Available Newport-branded “non-menthol” cigarettes were purchased in California in 2025, extracted and tested for sensory cooling activity by Ca^2+^ microfluorimetry of HEK293T cells expressing the human TRPM8 cold/menthol receptor. Chemical analysis was performed by gas chromatography - mass spectrometry (GCMS). “Non-menthol” and menthol cigarettes marketed in 2023-24 served as controls.

**Results:** Extracts from Newport-branded “non-menthol” cigarettes marketed in California in 2025 did not elicit sensory cooling activity. Chemical analysis confirmed the absence of menthol and any of the major commercial synthetic cooling agents.

**Conclusions:** The tobacco industry removed sensory cooling agents from “non-menthol” cigarettes marketed in California. However, this did not result in the market withdrawal of “non-menthol” cigarettes in the state. “Non-menthol” cigarettes in California continue to be marketed with package designs resembling those of former menthol cigarettes, signaling the potential presence of a characterising flavour.

## Introduction

Since December 21, 2022, the sale of most flavoured tobacco products has been prohibited in California.^1^ This ban includes menthol cigarettes that were previously favored by youth and young adults, women and non-Hispanic Black Americans.^2^ In the same month, tobacco companies introduced “non-menthol” cigarettes in California, advertising them with designs highly identical similar to former menthol cigarette brands, including R.J. Reynolds’ Newport (Figure 1A, B) and Camel and ITG Brands’ Kool cigarettes. ^3-5^ Total cigarette sales in California declined by 21.1% in the 18 months after the flavour ban, mostly due to the abrupt decline in menthol cigarette sales.^6^ However, this was partially offset by sales of the new “non-menthol” cigarettes.^6^ Chemical analytical and pharmacological studies revealed that some brands of “non-menthol” cigarettes introduced in December 2022, Newport Green and Camel Crisp, contained WS-3, a synthetic sensory cooling agent.^3 4^ California regulators determined that these cigarettes violated the State’s flavour ban; however, R.J. Reynolds filed a judicial complaint stating that WS-3-containing “non-menthol” cigarettes have no characterizing flavours.^7^ California lawmakers responded by revising tobacco legislation to include cooling sensations in the definition of characterizing flavours, thereby effectively banning WS-3-containing cigarettes.^78^

**Figure 1:**
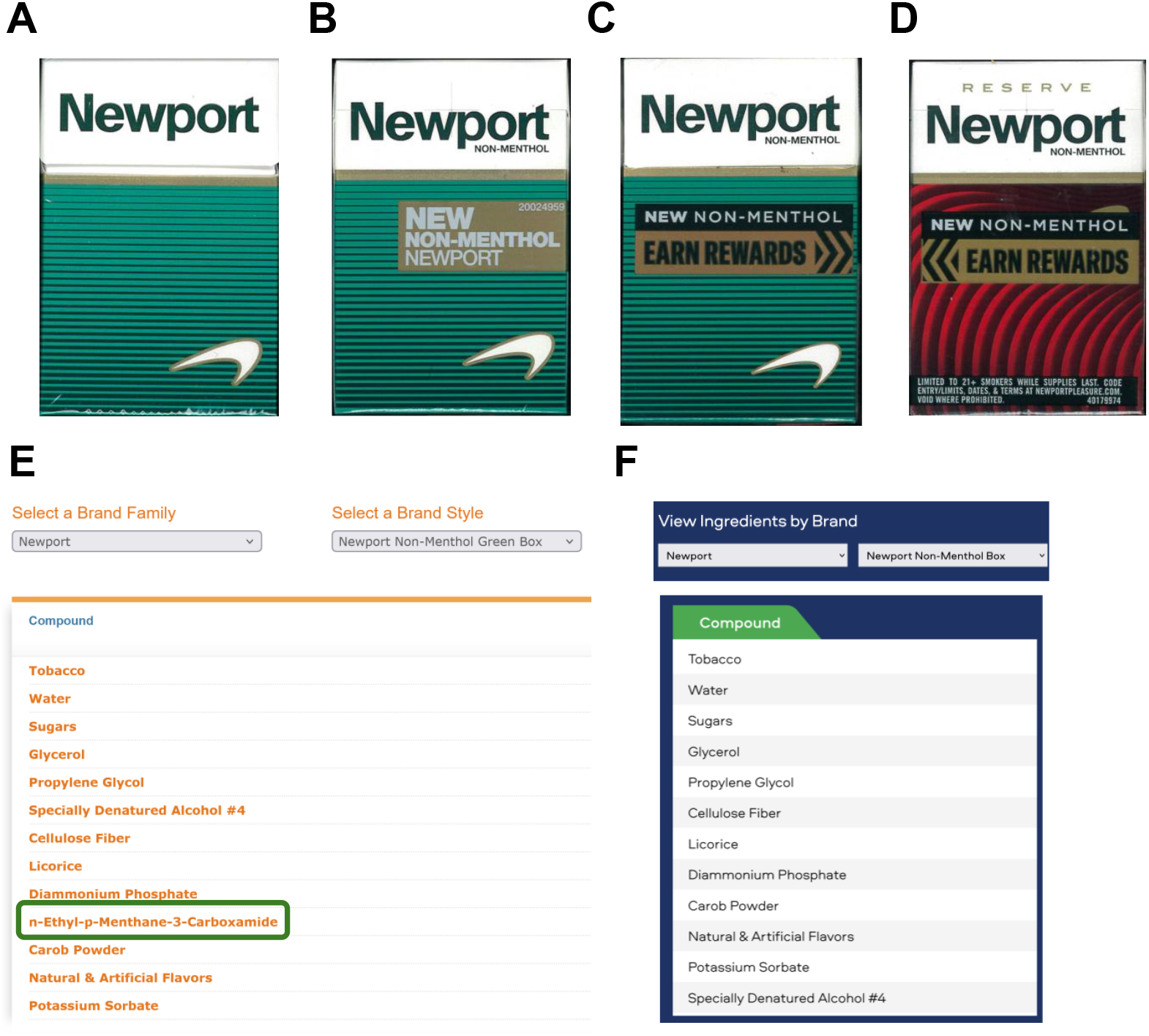
New “non-menthol” cigarettes introduced in California in 2025. **A** Newport menthol cigarettes box purchased in North Carolina in 2023. **B** Newport “Non-Menthol” green box purchased in California in 2023, in cellophane wrapper with white letters on gold background. **C** Newport “Non-Menthol” Box (green) purchased in California in March 2025, in cellophane wrapper with “New Non-Menthol” printed in white letters on black background, and “Earn Rewards” printed in black letters on gold background and with arrow pattern. **D** Newport “Non-Menthol” Reserve Box (red/black) purchased in California in March 2025. **E** Ingredient list of Newport Non-Menthol Green Box (shown in B), copied from the 2023 R.J. Reynolds commercial integrity website, with WS-3 (n-Ethyl-p-Menthane-3-Carboxamide) highlighted. **F** Ingredient list of 2025 Newport Non-Menthol Box (shown in C), copied from the 2025 R.J. Reynolds commercial integrity website.

In March 2025, after California’s new legislation took effect, two new “non-menthol” cigarette varieties were introduced by R.J. Reynolds, including i) a Newport variety with altered “new non-menthol” advertising on the wrapper, but otherwise in an unaltered green box, and ii) a Newport Reserve “new non-menthol” variety in a box with red / black wave patterns (Figure 1C, D). The company had also changed its “Commercial Integrity” website in which WS-3 was listed as an ingredient in its California-marketed Newport varieties introduced in 2022 (Figure 1E).^9^ On the revised website, WS-3 is not listed as an ingredient of the new varieties introduced in 2025 (Newport non-menthol Box, Newport Reserve non-menthol Box) (Figure 1F). However, the ingredient list includes “Natural & Artificial Flavors” which could include WS-3 or other synthetic cooling agents.^10^

## Methods

### Assessment of sensory cooling activity

Flavour chemicals were extracted from cigarette tobacco rods by stirring the rod material overnight in 10mL methanol. Extracts were strained from solid materials using a nylon strainer and subsequent centrifugation. Supernatants were collected and dried using vacuum centrifugation. Dried constituents were reconstituted in calcium assay buffer (Hank’s Balanced Salt Solution with 10 mM HEPES) and further dilutions were prepared (1X-200X) to assess sensory cooling activity by Ca^2+^ microfluorimetry of HEK293t cells expressing the human cold/menthol receptor, TRPM8 (FlexStation, Molecular Devices). 1X dilution is defined as the extract of one tobacco rod contents in 50 mL assay buffer, and 200X is 200-fold dilution thereof). Ca^2+^-responses from these extracts were normalized to the Ca^2+^-response elicited by a maximally activating concentration of agonist L-menthol (1 mM; TRPM8). Dose-response curves for receptor activity and associated calcium influx changes were plotted using non-linear regression analysis with a 4-parameter logistic equation (Graphpad Prism 9.0, San Diego, CA).

### Chemical analysis

Chemical analysis was carried out as described previously, ^3^ with slight modification: The tobacco filler was removed and placed in a 20 mL vial, and the wrapping paper and filter material were cut into smaller sections and also added to the same vial. 10 mL of methanol (Fisher Scientific, Waltham, MA) were added to the vial, and the vials were shaken using an orbital shaker (Thermo Scientifc Solaris, Waltham, MA) at 25 °C for 60 minutes. Extracts were filtered (0.22 µm, Millex-GP, Sigma-Aldrich, St. Louis, MO) and 1 _μ_L was injected directly into a gas chromatograph with connected mass spectrometer (GC/MS; Perkin-Elmer Clarus 580-SQ8S, Shelton, CT), which was outfitted with an Elite-5MS column (length 60 m, id 0.25 mm, 0.25 μm film, Perkin-Elmer) using the following heating program: Hold at 30 ºC for 7 min, ramp 10 ºC/min to 50 ºC and hold for 20 min, ramp 10 ºC/min to 310 ºC and hold for 7 min. Commercially available standards were purchased to confirm presence/absence of the following coolants: Menthol (Supplier: TCI America, Portland, OR; LOD: 3 µg/mL; LOQ: 9 µg/mL); WS-3 (TCI America; 5 µg/mL; 15 µg/mL); WS-23 (TCI America; 2 µg/mL; 6 µg/mL); WS-12 (Taima, Xi’An City, China; 50 µg/mL; 150 µg/mL); Frescolat MGA (as l-menthone glycerol ketal, Caldwell, NJ; 5 µg/mL; 15 µg/mL); Frescolat ML (as L-menthyl lactate, Sigma Aldrich; 10 µg/mL; 30 µg/mL); Frescolat X-Cool (Symrise, Teterboro, NJ; 10 µg/mL; 30 µg/mL).

## Results

To test whether the newly introduced Newport “non-menthol” varieties impart a cooling flavour, we purchased these cigarettes in March and May 2025 at convenience stores in the Alameda and San Francisco areas. Extracts from tobacco rods were serially diluted and superfused over HEK293T cells expressing the human TRPM8 cold/menthol receptor activated by menthol, WS-3 and other synthetic cooling agents, measuring activity by Ca^2+^ microfluorimetry as published.^3^ Extracts from Newport new “non-menthol” (green) and Newport new “non-menthol” Reserve cigarettes (red) did not increase intracellular Ca^2+^ levels, even at the lowest dilution tested (10x) (Figure 2). In contrast, control extracts of 2023 North Carolina-marketed Newport menthol cigarettes and 2023 California-marketed “non-menthol” green cigarettes increased intracellular Ca^2+^ levels in a dose-dependent manner, due to the verified presence of menthol or WS-3, respectively (Figure 2).^3^ These data suggest that the Newport “non-menthol” varieties introduced in California in 2025 contain neither menthol, nor WS-3, or any other sensory cooling agent at levels comparable to traditional menthol cigarettes, or the 2023 Newport “non-menthol” cigarettes.

**Figure 2:**
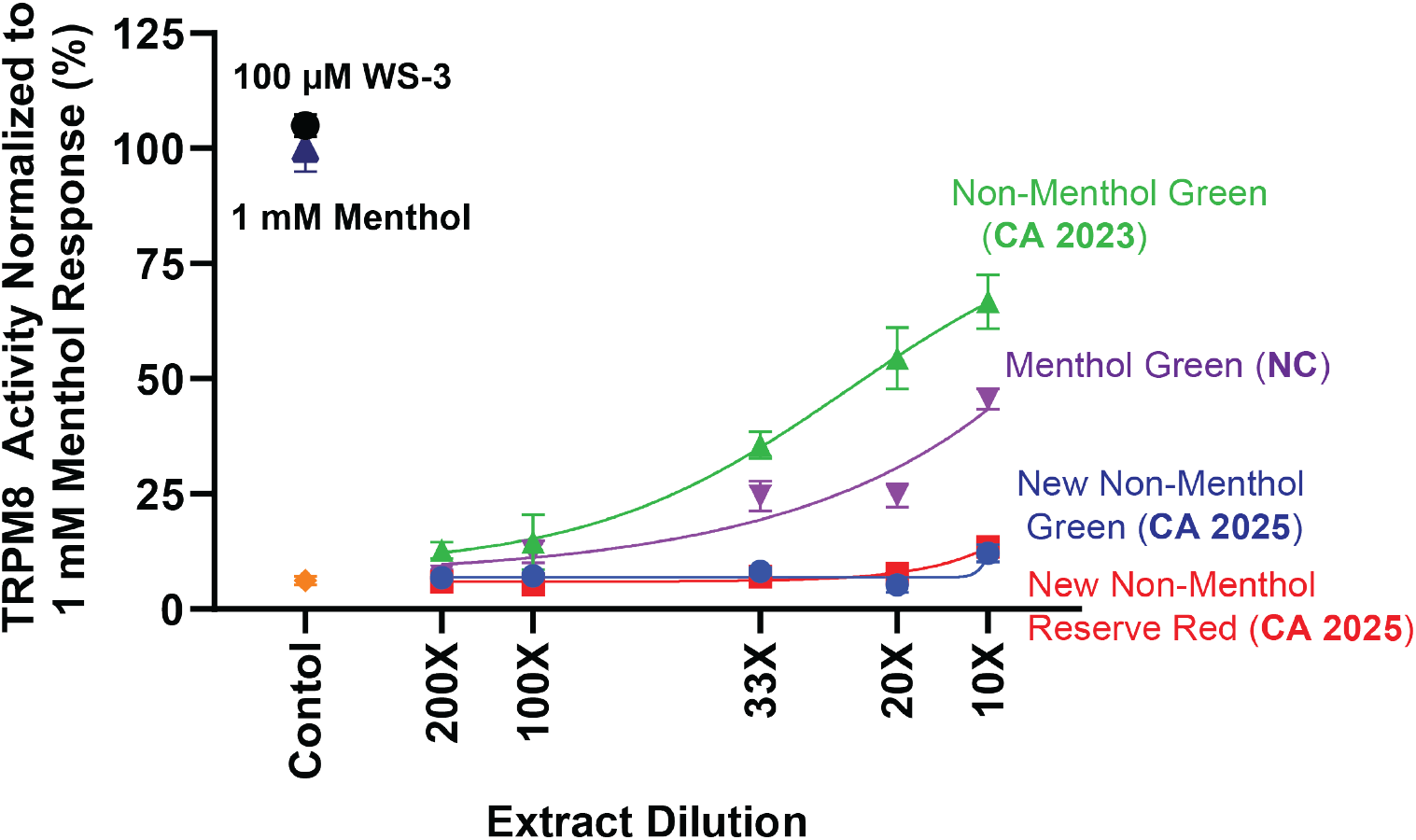
Sensory cooling activity of Newport “non-menthol” cigarettes introduced in California in 2023 and 2025, compared to Newport menthol cigarettes. Dose-response analysis of human TRPM8 cold/menthol receptor-mediated Ca^2+^-influx, upon superfusion of cells with dilution series of extracts from Newport brand cigarettes. Green: Non-menthol green box, purchased in CA in 2023; Purple: Menthol green box, purchased in North Carolina in 2023; Red: New non-menthol green box, purchased in CA in 2025; Black: New non-menthol reserve, purchased in CA in 2025. The increase in intracellular Ca^2+^, measured as fluorescence units (F_max_-F_0_), was normalized to the Ca^2+^-response elicited by a saturating concentration of agonist L-menthol (1 mM; solid black triangle). Response to a saturating concentration of WS-3 (100 µM; solid black circle) and vehicle control (orange diamond) shown for comparison. 1X dilution is defined as the extract of one tobacco rod contents in 50 mL assay buffer, and 10X is 10-fold dilution thereof.

To test for residual presence of cooling agents, whole cigarettes were extracted and analysed for the major commercial sensory cooling agents (menthol, WS-3, WS-23, WS-12, Frescolat MGA, Frescolat ML, Frescolat X-cool) by gas chromatography - mass spectrometry (GCMS) following published protocols.^3^ None of the Newport “non-menthol” varieties introduced in 2025 contained cooling agents over the limits of detection.

## Discussion

Taken together, these data demonstrate that R.J. Reynolds removed sensory cooling agents from their California-marketed Newport “non-menthol” varieties. However, it is important to note that in the case of the Newport non-menthol green variety (Figure 2C), the box design and color is identical to the previous WS-3-containing version, and very similar to Newport menthol cigarettes (except the “non-menthol” label), which could continue to signal the presence of a minty and cooling characterizing flavour to people who previously smoked menthol cigarettes. R.J. Reynolds’ strategy is similar to ITG Brands’ for their Kool “non-menthol” cigarettes introduced in California in December 2022 that have blue/black or green/black box coloring similar to the brand’s menthol cigarettes, but do not contain any cooling agent.^3 4^ A distinctive choice of package colouring and design by the industry is a common strategy to manipulate consumer expectations about the flavour and strength of tobacco products.^11^ In June 2023, California regulators raised concerns about the misleading package design and coloring of these Kool “non-menthol” cigarettes. ^12^ Similar concerns should be raised about the new Newport non-menthol green variety introduced in 2025 that continues to signal that a characterising flavour could be present.

## Declarations

### Disclaimer

The funding organizations had no role in the design and conduct of the study; the collection, management, analysis, and interpretation of the data; the preparation, review, or approval of the manuscript; nor in the decision to submit the manuscript for publication. The content is solely the responsibility of the authors and does not necessarily represent the views of National Institutes of Health or the Food and Drug Administration.

## Author Contributions

SVJ, HCE and SEJ conceptualized and designed the study; SKS and SO provided advice on product choice and regulatory policy. SVJ acquired, analyzed and interpreted the calcium microfluorimetry and receptor activity experiments, supervised by SEJ; HCE acquired and analyzed the chemical analytical data, supervised by PTA and JBZ. SVJ, SEJ and HCE drafted the manuscript; All authors critically reviewed, edited, and approved the final manuscript. SEJ attests that all listed authors meet authorship criteria and that no others meeting the criteria have been omitted.

